# DEED: A Dataset for Dream-related Emotion Research

**DOI:** 10.1101/2022.09.19.508475

**Authors:** Wanqiu Liu, Yupeng Zhang, Pengcheng Ma, Lei Zheng, Di Zhou, Zhengbo Chen, Mingming Shen, Yongchun Cai, Zhengyi Lu, Qiao Liu, Junwen Luo, Xiaoan Wang

## Abstract

Emotion is closely related to human cognition and behaviour. In recent years, scholars have conducted extensive research on emotion in waking state based on electroencephalography (EEG) and achieved certain results. However, Emotional activity continues after sleep, with a concentrated response of sleep emotions in dreams. Sleep emotions are concentrated in dreams, which can better reflect a series of real physical and psychological states of the human body. Currently, there is no publicly available dataset for the assessment of dream mood. Therefore, we present a physiological dataset Dream Emotion Evaluation Dataset (DEED) for the assessment of dream mood, which recorded EEG signals from 38 participants over 89 whole sleep nights and 533 dream segments(after exclusion of unqualified nights, those dream segments are extracted from 82 whole sleep nights). We studied the correlations between the subjective ratings and the EEG signals and brain network patterns for dream emotions. In addition, the relationship between the asymmetry of left and right brain bands and positive and negative dream emotions was studied. The machine learning algorithm was also used to classify different emotional EEG, which confirmed the validity of the dataset. In the meantime, we encourage other researchers to explore the underlying neural mechanisms involved in sleep.

## 1 Introduction

Emotion is a psychological and physiological response caused by various external stimuli[1]. Closely related to human cognition and behaviour, it is a comprehensive reflection of the physiological state of various everyday thoughts and behaviour and plays an important role in people’s daily lives. In recent years, research around emotions has been applied in psychology, neurobiology, medicine, and other related fields with some results[2,3]. At present, research on emotions includes external human responses such as facial expressions, verbal expressions and gesture changes, as well as internal human responses including electroencephalography (EEG), electrocardiogram (ECG) and magnetoencephalography (MEG)[4]. Meanwhile, research on related emotions based on neural mechanisms has also started at a very early stage. EEG has timeliness, can reflect the real-time brain function changes of subjects, and has the characteristics of objectivity, less prone to artifacts, low measurement cost[5], and so it has been widely applied in the field of emotion recognition.

In studies related to EEG-based neurophysiology and emotion processing, researchers have identified a type of local EEG activity associated with emotion in the prefrontal cortex of the brain. This activity arises from changes in neural activity associated with this brain region and correlates with current emotional patterns[6], called EEG frontal alpha asymmetry (FAA). FAA is widely used as a biomarker for exploring mood, anxiety, insomnia, and traumatic stress syndromes in various types of psychopathology, cognitive regulation[7], and other fields.FAA responds to changes in the current emotional state or in the functional state of recognition behaviour[8] and numerically reflects the difference between the alpha power (8-13 Hz) in the right and left hemispheric prefrontal cortex, usually calculated based on F4-F3 leads. Due to the mechanism of the contralateral inhibitory connections between the two hemispheres, higher FAA scores are reflected when there is an elevated activity in the right prefrontal or reduced activity in the left. When right-sided EEG activity is stronger, it is usually associated with withdrawn emotional activity, while more left-sided prefrontal activity is associated with approach emotional activity. In addition, studies had shown that EEG in beta and gamma bands are also used to identify happy and sad emotions when the body is in different evoked emotional states[9]. With the discovery of various markers of EEG differences between emotions, various computer-based algorithms for automatic emotion recognition based on EEG have been widely developed.

EEG has many non-linear features and is susceptible to interference, so various features need to be extracted to focus on the emotions at this time[10]. Generally, statistical parameters, non-stationary indices, Hjorth parameters, etc. can be extracted based on the time domain of the EEG; power spectrum density (PSD), differential entropy (DE), fractal dimension(FD), etc. can also produce effective features for identifying emotions. Smith[11] proposed an optimised variational mode decomposition using single-channel EEG, selecting the dominant channel by the eigenvector centroid method, and extracting the Hjorth complexity, differential absolute standard deviation value (DASDV), log-energy entropy (LEE), and Renyi entropy (RE), by using machine learning classifiers such as random forest(RF), support vector machine(SVM), and k-Nearest Neighbour(kNN), the EEGs of four emotions: fear, happy, sad, and relaxed were identified, obtaining an average accuracy of over 90%. Edgar[12] used audio and video to induce emotion generation by extracting DE, differential asymmetry, and rational asymmetry symmetry, after that used a mutual information matrix for feature selection to identify the four emotions using algorithms such as RF, obtaining an accuracy of 82.51%. More and more deep learning-based emotion recognition models have been proposed with the public availability of some evoked emotion EEG datasets (e.g. SEED, DEAP, DREAMER, AMIGO, etc.). Since the two hemispheres have different EEG activity in different emotional states, based on the hemispheric asymmetry, Yang[13] designed a model called bi-hemisphere domain adversarial neural network (BiDANN) to implement EEG recognition of different emotions and tested the performance of the model on the SEED dataset and achieved accuracy over 92% in recognising the three emotions.

Emotional activity continues after sleep, with a concentrated response of sleep emotions in dreams. Nathaniel[14] first showed that dreaming occurred during a specific sleep phase that was accompanied by higher levels of cortical EEG activity and rapid eye movements(REM), which provided a neurophysiological marker for the emergence of dreams. As scholars have studied dreaming more closely, some subjects who wake up during non-rapid eye movements(NREM) periods also report dream production[15]. However, subjects who are awakened during the REM period, over 70% recall the content of their dreams, while a smaller proportion of subjects recall the content of their dreams during the NREM period, and subjects who are awakened during REM tend to report more vivid and complex emotional experiences. Some studies have shown that this stage of sleep is often accompanied by the consolidation of emotional memories[16], so the current research on dreams is mainly from the REM period.

Research has shown that there is a degree of correlation between the content of dreams and waking life experiences in humans[17], with dream characteristics reflecting waking life experiences and dreams influencing human thought activity to some extent. The dream experience can take many forms[18], including not only simple perceptions and images but also thoughts and experiences of various things, similar to during the waking state. Some studies have also shown that when the human body is dreaming, the subjective feelings generated in dreams have a similar neural coherence to the waking state, such as thinking, perceiving, acting, and thinking in dreams. Therefore, the emotions generated in dreams can share the same neural mechanism as those generated in the waking state[19]. From a physiological perspective, emotions in sobriety are more likely to be influenced by external environmental factors, and when asleep, its state is largely disconnected from the external environment, and the emotions are more reflective of the body’s physiological and psychological state. Hence, the study of dream emotions has the same importance as the study of the waking state.

Currently, the most valid measure of dream emotion is the verbal report after waking, however, most of the dream content is forgotten after a full night’s sleep, so the most reliable dream emotion can only be recorded if the subject is awakened for a verbal report immediately after the REM period. For these reasons, research on dream emotions is sparse and there are no published datasets relating to dream emotions. Our search revealed only a few studies of emotions based on the REM period EEG. Daoust[20] investigated the relationship between dream emotions and EEG-related properties in normal and autistic individuals; the results suggest that the alpha activity of dream EEG during the REM period may represent the neurophysiological basis for the association with dream emotions. Sikka[21] designed an experiment in which subjects were awakened five minutes after the onset of a rapid REM period and asked about the type of dream emotion that the subject felt. The EEGs during dreaming were collected and FAA was calculated for channels F3-F4. The results showed that the FAA could be used to predict the anger index in dreams during REM sleep as well as during the waking state in sleep. In contrast, research on computer-automated algorithms for the recognition of emotions in REM dreams has not been reviewed in the literature.

Therefore, we designed an experiment for acquiring different dream emotion EEGs during the REM period, which were organized into the DEED and released publicly. DEED contributions include, but are not limited to:

- A dataset of larger samples with non-stationary characters, obscure spatiotemporal features, and time-series signals encoded with dream mood information. This is suitable for testing algorithms with biological-like intelligence performances such as few-shot learning, generalization, and anti-interference ability.
- It enables potential exploration of the biology of dreams, the mechanisms of sleep-emotion interactions, and the neural manifestations of the transition between consciousness and unconsciousness.

In this paper, our first aim is to publicly present the DEED dataset, explore the correlation between EEG and subjective ratings, perform a clustering analysis of brain network patterns, and investigate the correlation between left and right brain asymmetry indices and positive and negative dream emotions in each waveform. Another aim is to validate the effectiveness of machine learning algorithms such as SVM, kNN, LightGBM and GBDT and RF in classifying the EEG of REM dreams under different emotions by extracting features.

## 2 Materials

### 2.1 Participants

This experiment was conducted among current students at Zhejiang University and a total of 38 subjects(18 females, 20 males, age 21.16 ± 1.81) volunteered to participate in this experiment (recruitment was done through emails sent by current students). Participants were screened through the following questionnaires: (1) healthy, no brain disease; (2) right-handed, no physical disability; (3) native Chinese speaker; (3) good sleep quality(Pittsburgh Sleepiness Scale, Epworth Sleepiness Scale score ≤ 16); (4) good mental status, no depression or other psychiatric disorders(DSM-5 Self-rated Level 1 Cross-Cutting Symptom, screening criteria: score ≤ 8; State-Trait Anxiety Inventory (STAI), screening criteria: score ≤ 39); (5) not using medication. The described study was reviewed by the Research Ethics Committee of Zhejiang University and informed written consent was obtained from all participants. All participants were paid accordingly.

### 2.2 Experiment design

We design a new emotion experiment to collect EEG data, which are different from those other existing publicly available datasets. The screening questionnaire was completed three days before the start of the experiment. The subject who met the entry criteria for data collection was required to arrive at the sleep room at around 20:00 and to wear the Polysomnography(PSG, Alice 6 LDE) under the guidance of the experimenter, with the electrodes positioned as shown in Figure 1. After completing the Karolinska Sleepiness Scale, the subject was asked to lie supine to collect the resting EEG for both four minutes each eyes opened and eyes closed, and then to watch one of the screened films with positive, negative and neutral emotions, with each of the three emotion categories being viewed during the three sessions. After watching the film, the subject was given the Positive and Negative Emotion Scale(PANAS) and the resting EEG has then collected again for four minutes with both eyes opened and eyes closed. The subject was reminded to lie as flat as possible and turned off their mobile phones and other electronic devices and wore earplugs to protect them from outside distractions.

**Figure 1.**
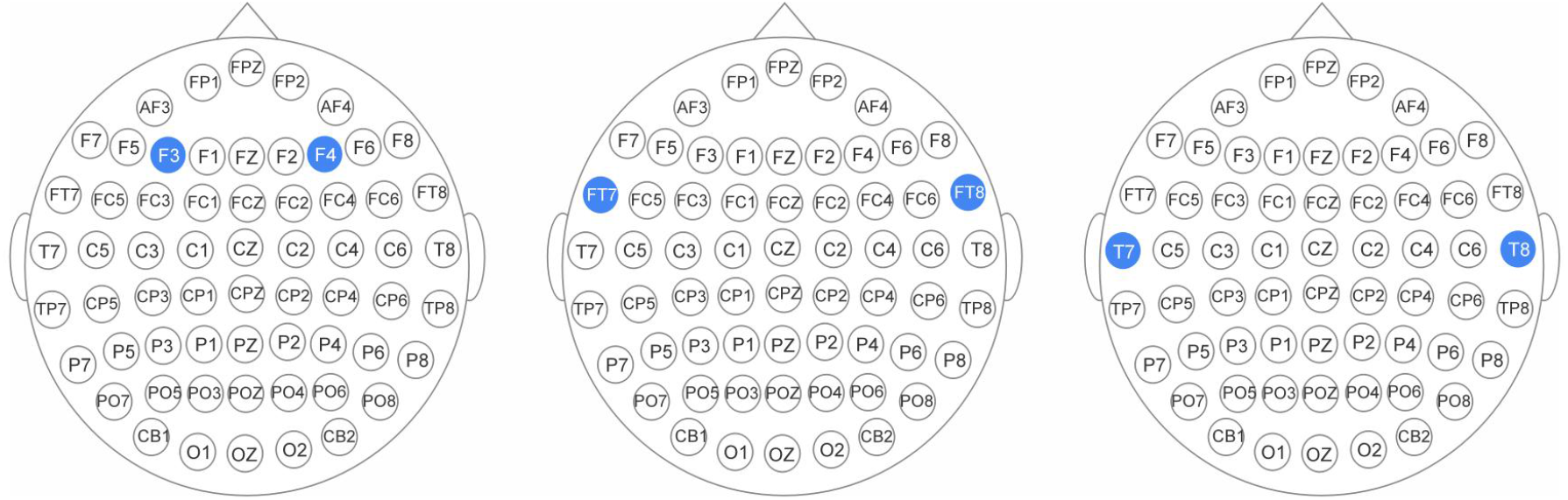
PSG electrode position

The subject’s EEG was continuously observed and monitored by the experimenter after waking up at 5:00 a.m. Whenever the REM phase was monitored to occur, the subject was awakened using the tone signal which lasted for five minutes [21] and upon waking up, the subject provided a verbal report according to the task, including what he or she was thinking at the last moment before waking up, what his or her mood was at the moment, and the experimenter confirmed with the subject the current moods were recorded and included five categories: very unhappy, relatively unhappy, calm, happy and very happy, skipping this step if no dreams occurred. The experimenter then allows the subject to continue sleeping and repeats the previous steps when the REM phase is detected again. This process continued until the subject fully awoke. The process is shown in Figure 2.

**Figure 2.**
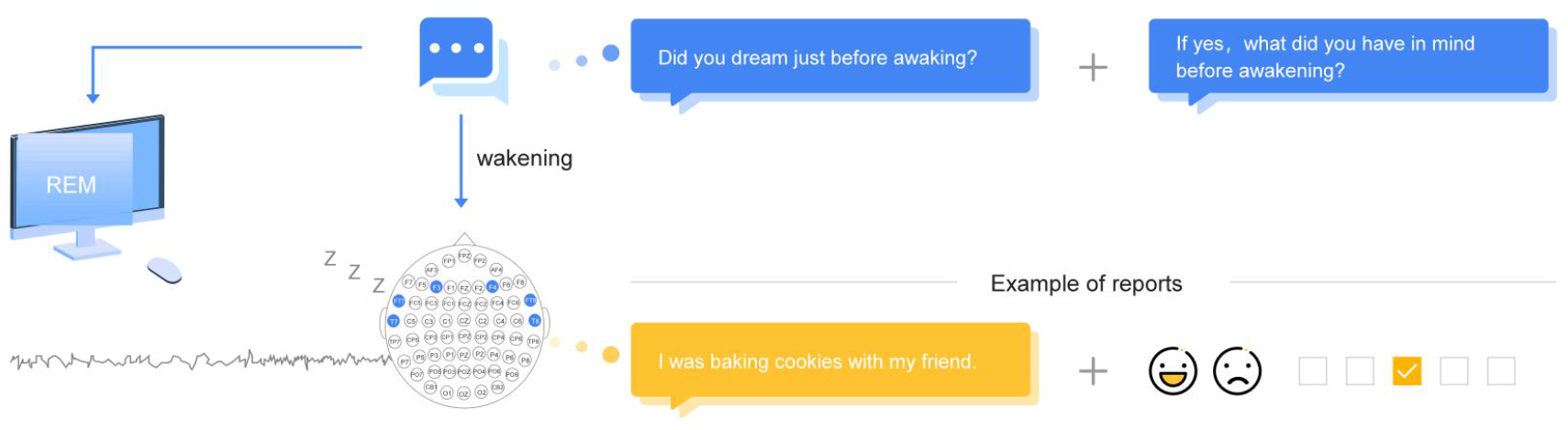
The structured interview during the REM period. Master(small dialogue box symbol) observes the brain waveform of the subject’s sleeping state through the screen. Once they detected the appearance of the REM phase, they woke up the subject(electrode position brain symbol) and asked the subject to answer a series of questions designed in advance.

After the last REM period was reported, the subject was asked to lie flat on the bed and the resting EEG was collected for four minutes each in the eyes-opened and eyes-closed state, similar to before falling asleep, after which the PSG device was removed to end the experiment. At the end of the experiment, the subject was allowed to continue resting in the sleep room until 12:00. The overall experimental flow is shown in Figure 3. In this paper, data from yellow segments collected during the early morning stages of rapid eye movement sleep are processed and analysed.

**Figure 3.**
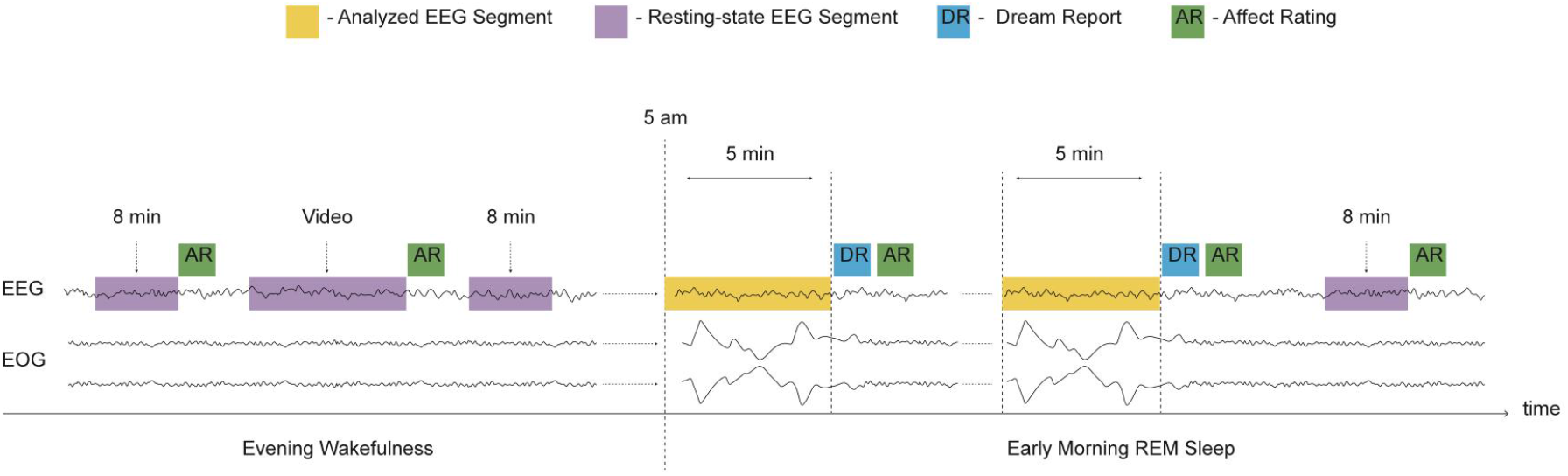
Experiment design Evening Wakefuless: Experimental flow diagram. After completion of PSG device wear, subjects’ EEG were recorded for eight minutes in the waking state, followed by a recording of their current waking emotion (AR). Emotion elicitation was then performed by watching a movie and recording one waking emotion, and after completing a second eight minutes waking state resting EEG the subject was allowed to fall asleep. Early Morning REM Sleep: After 5 am, REM sleep acquisition was performed and participants were awakened using tone signals five minutes after the onset of REM. After each awakening, the subject was asked about their mood and a dream report (DR) was generated, after which the subject was allowed to continue sleeping and the process continued until fully awake. An 8minutes awake state resting EEG acquisition was then performed, after which the experiment ended.

### 2.3 Data acquisition and Description

Six of the gold cup electrodes (F3, F4, FT7, FT8, T7 and T8) were placed on the participants’ scalps according to the 10-20 system. Two electrodes were used to record EOG (superior to the left eye and inferior to the right eye). One electrode placed on the chin was applied to record EMG. The ground electrode was placed on the forehead. All electrodes, except the bipolar EOG and EMG electrodes, were referenced to the right mastoid (M2). The electrode placed on the left mastoid (M1) was also used to record data for re-reference. All impedance was kept below 20kΩ.

The EEG data were first resampled at 200Hz, then notch-filtered at 50Hz. Then they were bandpass filtered using 1 Hz and 45 Hz cutoff values. By following rules in AASM (American Academy of Sleep Medicine), each REM was tagged and sliced. Furthermore, data recorded by each channel was re-referenced: data recorded from the left electrodes was subtracted by (M1-M2)/2, while the right was added by (M1-M2)/2 [22]. Additionally, AMICA in EEGLAB toolbox (version 2021.1) was applied to remove artifacts.

The extracted REM-phase data was then mood-labelled, and if the number of REMs for a subject in a particular experiment was less than four, we considered that night to have produced poor sleep quality in the subject and the results of that experiment were excluded from our analysis. After screening, valid data were collected for a total of 82 nights from 38 subjects.

#### 2.3.1 Raw EEG Data with Whole Night

The raw data for DEED is available at http://www.deeddataset.com//download. The 89 sleep files (.mat) recorded signals during an overnight sleep study contain Electroencephalogram (EEG) to identify sleep stages.

This raw EEG data include 7 EEG channels, and the channel number and the channel content as Table 1 shows, which the M1-M2 channel is used for reference.

**Table 1.**
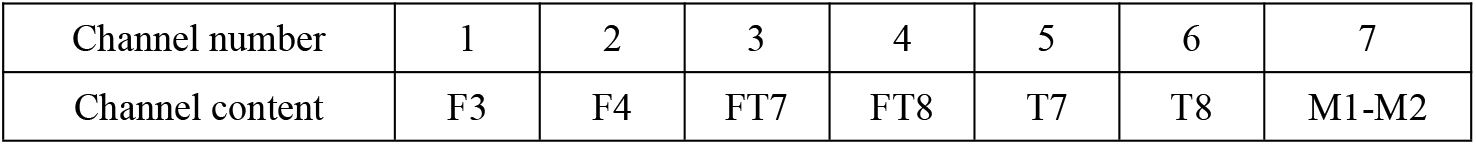
Channel number and channel content

#### 2.3.2 Dream EEG with emotion labels

EEG with emotion labels contains data that has been intercepted, organized and extracted from the raw data. In total, there are five hundred and 533 clips with their emotional label. They are all provided in the MAT format.

These files are named in a certain way: EEG_{subject ID+number order of the experiment}_{Movie}_{Emotion}_{R+order of REM}_{sleep stage before REM}. For example, EEG_S0021_M1_E3_R01_N2 represented that in subject S002’s first experiment, he/she watched a negative movie(1 represented negative, 2 represented neural and 3 represent positive), awakened in the first REM period, was in N2 stage sleep before waking and reported this dream was neutral mood (E0 represented no dream, E1/E2 represented negative mood, E3 represented neutral mood, and E4/E5 represented positive mood). To save the researcher from having to repeat the pre-processing and file organization work, each file inside this zip is downsampled, filtered, re-referenced, and sliced.

After pre-processing, a total of 533 EEG data was extracted during the REM period, of which a total of 153 were labelled as positive emotions, 96 were labelled as negative emotions, 244 were neutral emotions, and the remaining 40 of which had no dreams.

## 3 Analysis of the correlates between EEG and subjective ratings

To investigate whether brain activities show different patterns in different dream emotional contents, we calculated the correlations between the subjective ratings and the EEG signals. The frequency power of each trial between 1 and 45 Hz was extracted with Welch’s method with windows of 200 samples (1 second). These changes of power were averaged across 6 electrodes over the frequency bands of delta (1-4 Hz), theta (4-8 Hz), alpha (8-14 Hz), beta (14-32 Hz), and gamma (32-45 Hz). For the correlation statistic, we computed the Pearson correlation coefficients between the power changes and the subjective ratings. And a permutation test (Bonferroni) is used to determine the significance of Pearson correlation coefficients, where we random the procedure 200 times to mismatch the power and the corresponding subjective rating to calculate the chance level for the correlation. The detailed correlation between the subjective ratings and the power changes is shown in Table 2. According to the permutation test, the correlation between the subjective ratings and the power changes was higher than the chance level in delta, theta, alpha, and beta bands (P < 0.05). And the correlation in the alpha band is significantly higher than the chance level (P < 0.01). It indicates the importance of alpha oscillations for investigating emotion during dreams, which is consistent with previous research [23]. However, the correlation in the gamma band shows lower than the chance level. The reason may be caused by the amplitude in gamma bands which is lower than in other frequency bands. The power in gamma bands is more easily affected by noise than in other frequency bands. Therefore, external noise attenuates the correlation effect in it.

**Table 2.**
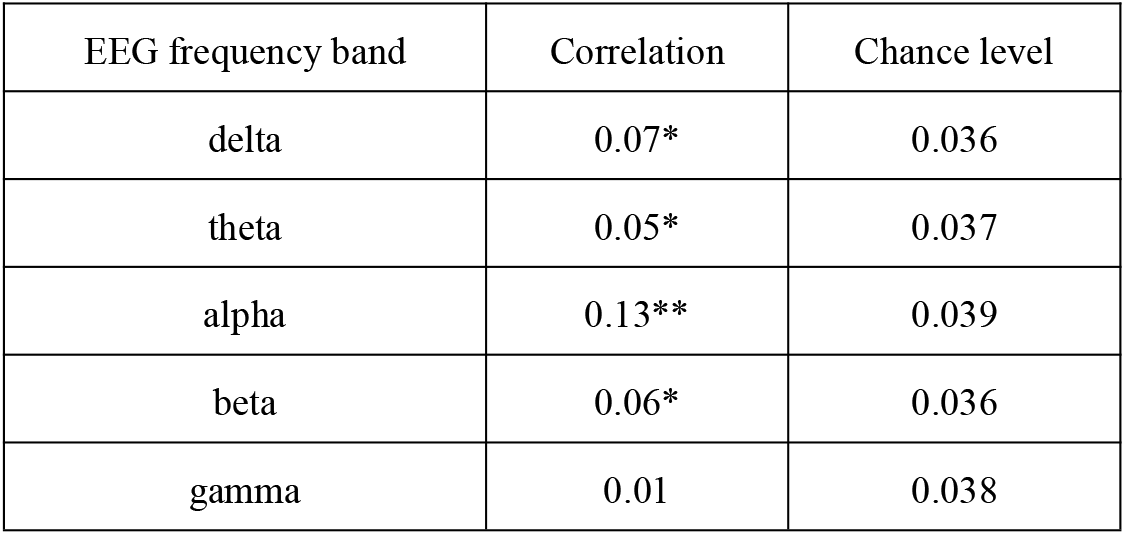
Correlation between the subjective ratings and the power changes (* P < 0.05; ** P < 0.01)

## 4 Brain network patterns and dream emotions

Cognitive processes are supported by the transient coupling and uncoupling of activity between distant brain regions [24]. The interactions between brain regions can be modelled as a functional brain network, with nodes and edges representing the brain regions and functional connectivity between brain regions, respectively [25]. To address whether the different dream emotions have specific brain network patterns, this study uses the six nodes (six electrodes) and Pearson correlation to describe the functional link among the nodes. This would result in 15 edges if linking all pairwise nodes for EEG each trial. Finally, these parameters of the brain network in each trial were clustered by using the unsupervised k-means algorithm. According to the 5-point rating scales from participants during the experiment, all the REM clips were divided into three classes (negative, neutral, and positive). And the threshold was simply placed in the middle of the 5 points rating scales (1-2: negative; 3: neutral; 4-5: positive). Therefore, we set the cluster number to 3, and we performed 1,000 k-means repetitions with random initial states. By calculating the corresponding F1 scores (the harmonic mean of precision and recall) between the actual grouping and the result of the clustering, this study investigates the relationship between different dream emotions and brain network patterns. The final cluster result is shown in Table 3. The chance level is also calculated from a permutation test (Bonferroni), where we random the emotion label 200 times to get the cluster results. According to the F1-score of K-means, we found the unsupervised cluster results show a significant difference with chancel level at the whole frequency bands. The results may indicate the different brain network patterns for dream emotions.

**Table 3.** Results of F1-score for K-means based on the brain network pattern (** P < 0.01)

| EEG frequency band | F1 score | Chance level |
| --- | --- | --- |
| delta | 0.40** | 0.33 |
| theta | 0.38** | 0.33 |
| alpha | 0.41** | 0.34 |
| beta | 0.43** | 0.33 |
| gamma | 0.46** | 0.33 |

## 5 Hemispheric Asymmetry Analysis

Short-time Fourier changes were performed on data segments from each preprocessed completed REM period using a 0.5 overlapping hamming window to obtain the mean spectral power of the signal. The data were then normalized by the natural logarithmic transformation of the d delta, theta, alpha, beta, and gamma band powers at the prefrontal (F3-F4), front-temporal (FT7-FT8), and temporal (T7-T8) electrodes. The scores for the left and right brain asymmetry indices were then calculated by subtracting the power of the left hemisphere electrode (ln right - ln left) from the power of the homologous right hemisphere electrode. Based on SPSS (version 20, IBM) software, the Wilcoxon Signed-Rank test was applied to the left and right brain asymmetry indices in the delta, theta, alpha, beta and gamma bands of leads F3-F4, FT7-FT8, and T7-T8. The test was used as a statistical analysis of positive and negative emotions, setting a confidence interval of 0.95, that is, when P<0.05, the difference in asymmetry indices between positive and negative emotions was considered statistically significant.

By analyzing the left-right symmetry indices in each band of the F3-F4 leads, we found statistically significant differences in the delta (P=0.008, Z=-2.440), theta (P=0.002, Z=-3.088), alpha (P=0.007, Z=-2.711), beta (P=0.017, Z=-2.397) bands. However, the gamma (P=0.263, Z=-1.118) band was not statistically significant. By analyzing the left-right symmetry indices in each band of the FT7-FT8 leads, we found statistically significant differences in the delta (P=0.049, Z=-1.944), alpha (P=0.007, -2.686), beta (P=0.034, -2.101), gamma (P=0.001, Z=-3.819) bands, but the theta (P=0.140, Z=-1.474) band was not statistically significant. Meanwhile, by analyzing the left-right symmetry indices in each band of the T7-T8 leads, we found statistically significant differences in the theta (P=0.028, Z=-2.193), alpha (P=0.042, Z=-1.864), beta (P=0.033, Z=-2.138), gamma (P=0.004, Z=-2.770) bands. However, the delta (P=0.217, Z=-1.235) band was not statistically significant.

We calculated hemispheric asymmetry indices across the frequency bands for both positive and negative emotion groups. The positive values represent stronger right-side brain activity and weaker left-side brain activity, and the larger the absolute value, the greater the difference in brain activity between the two sides. A histogram of the asymmetry index for each wave of the different emotions provides a visual representation of the changes in this index, and the results are shown in Figure 4.

**Figure 4.**
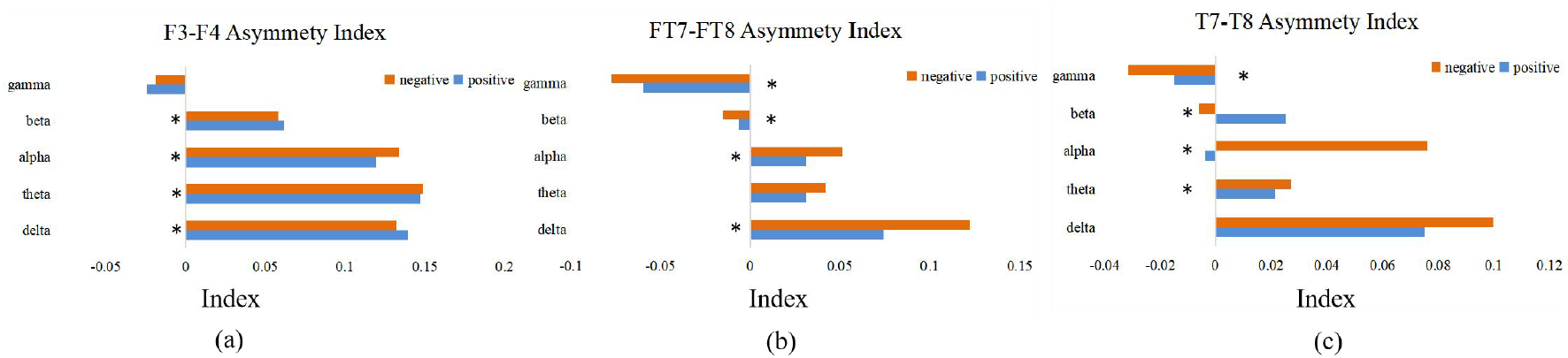
Left and right brain asymmetry indices. (a) shows the distribution of hemispheric asymmetry indices between positive and negative emotions in each band of the frontal cortex (b) shows the front-temporal hemispheric asymmetry indices (c) temporal hemispheric asymmetry indices. The horizontal axis represents the specific value of the asymmetry index, the vertical axis represents the different frequency bands of the EEG, and ∗ represents a statistically significant difference in the asymmetry index between positive and negative emotions

We found that when subjects experienced dreams with positive emotions, their frontal symmetry index were larger than that of negative emotions in the delta, beta bands, and the EEG activity in the right hemisphere were stronger than in the left hemisphere. The EEG activity in the right hemisphere was generally weaker in the theta, alpha bands than in the left hemisphere. Analysis of front-temporal and temporal cortex asymmetry, we found there are differences in the left and right hemisphere asymmetry index analysis between positive and negative emotions.

## 6 Dream Emotion Classification

In this chapter, to evaluate the quality of our data and baseline classifier, we performed binary classification and multi classification on REM period data. The purpose of binary classification is to detect negative and positive dream contents from EEG data. Meanwhile, a multi classification task is proposed for distinguishing the negative, neutral and positive dream contents.

Through the summary of the results of previous scholars’ research based on the recognition of waking state emotion EEG, we selected three features that are more widely used and have good performance: PSD, DE features, and CSP [26][27][28]. Their performance in recognizing dream emotions was tested in five classical machine learning classifiers:kNN, SVM(RBF), LightGBM, GBDT, and RF.

### 6.1 Feature Extraction

PSD describes the variation of the power of a signal with frequency. In this paper, we use Matlab’s FFT function to decompose the EEG, then take the mode of the complex exponential signal, and calculate the logarithmic sum after taking the one-sided spectrum to obtain the feature of the EEG in this frequency band. The average PSD was extracted by Welch’s method with a window size of 200 samples (1s) for each channel. As a result, we got a size of 30 (channel×frequency bands) PSD feature matrices from EEG data for each trial.

DE is a generalised form of Shannon’s entropy of information on continuous variables:

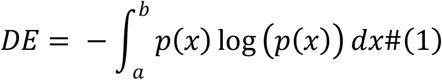

Where *p*(*x*) notes the probability density function of continuous information and [*a, b*] notes the interval over which the information takes its value. The DE of an EEG of a particular length, which approximately follows a Gaussian distribution 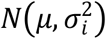 is:

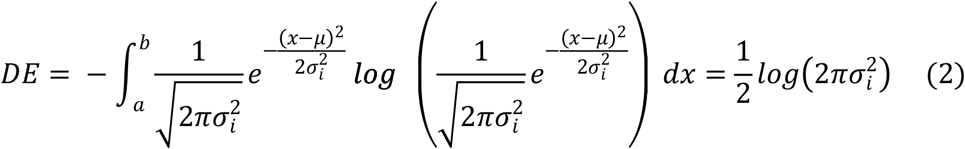

equal to the logarithm of its energy spectrum at a particular frequency band. And the average DE for each channel was calculated according to previous research [29] with a window size of 200 samples (1s). As a result, we got a size of 30 (channel×frequency bands) DE feature matrices from EEG data for each trial.

CSP algorithm belongs to the Spatial-Frequency domain feature and is often applied to binary classification problems under multiple channels. The co-space mode is a matrix of the number of channels multiplied by the number of samples put into a space for two types of emotional signals. Then, using the diagonalization of this matrix, we achieve a set of optimal spatial filters for projection, so that the difference in the variance values of the two types of signals is maximised, to obtain a feature vector with a high degree of differentiation[30], used in the next step to feed the feature vector into the classifier for classification. The operation process is as follows. Suppose there are n segments of EEG classified into two classes each of which is a matrix of the number of channels∗sampling points, and then these signals are taken as the covariance matrix Σ^+^ and Σ^−^, for the eigenvalue decomposition Σ^+^ + Σ^−^ =UDU^T^, where *U* is the eigenvector matrix and *D* is the diagonal matrix of eigenvalues, then the whitening value matrix can be found as 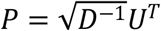, and then the whitening of Σ^+^and Σ^−^can be obtained as:

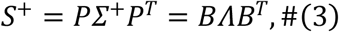

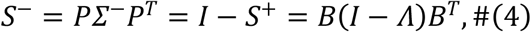

where *B* is the eigenvector matrix and is the matrix of eigenvalues. Then there is the projection matrix *W* =*B*^*T*^*P* as an eigenmatrix, as *B*^*T*^*P*Σ^+/-^*P*^*T*^*B=B*^*T*^*S*^+/-^*B*, each row in *W* as a vector is a copy of the spatial filter, and the larger the eigenvalue corresponding to its row vector, the larger the variance value difference. The row vectors can be sorted according to the corresponding eigenvalue size after taking the previous *n* row vector to construct the n-dimensional co-space feature can be, in this paper we find the best *n* = 4.As the result, we finally get the CSP features of size 20 (n-dimensional × frequency bands) from four CSP components in each frequency band.

### 6.2 Classifiers

KNN, SVM and RF are commonly used classifiers in machine learning and can be used as baseline classifiers in EEG-based emotion classification studies [11].

GBDT is an integrated learning algorithm constructed using the Boosting method[31]. The main idea is to iteratively train a series of weak classifiers, using a gradient boosting tree to accumulate the fitted residuals each time. The weak classifiers are CART regression trees. In a binary classification problem, the following formula is generally used as the loss function:

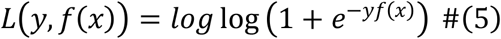

*y* ∈(-1,1)

where *f*(*x*) is the predicted result of the sub-classified, which is the fitted residual obtained based on the result of the previous iteration. *y* is the label of the data. The algorithm is to find the weak classifiers in the next iteration to minimise the loss function, and to sum up the results of each weak classifier to obtain the final result. Its *f*(*x*) initialisation function is shown as follows:

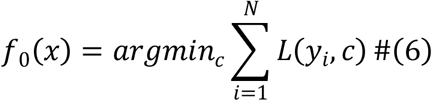

where the value *c* is the mean of the label values of all training samples and is the number of samples.

LightGBM uses a leaf-wise growth strategy to construct the number of trees. From all the current leaves, find the one with the largest splitting gain, and then split it, and so on. With the same number of splits, leaf-wise growth can reduce errors and increase accuracy. To improve the efficiency of GBDT, lightGBM has adopted two solutions[32]. The first one is the GOSS (Gradient-based One-Side Sampling) algorithm, which downsamples the samples by keeping those sample points with large gradients, while randomly sampling the sample points with small gradients proportionally. The second is the Exclusive Feature Bundling(EFB) algorithm, which constructs a weighted, undirected graph with vertices as features and edges as weights. Each weight is related to the conflict between the two features connected by the weight. The weights are then sorted in descending order, then assigning the features with greater conflict to existing feature bundles or creating a new feature bundle to minimise the overall conflict.

The histogram algorithm[33] is used to reduce the computational effort when calculating feature conflicts by first discretizing a continuous floating-point feature value into an integer, and then constructing a histogram of width k. After traversing the data once, the histogram accumulates the required statistics and is then traversed to find the optimal partition point based on the discrete values of the histogram. After a histogram has been constructed, histogram subtraction can be used to generate histograms for neighbouring leaves at a very small cost.

### 6.3 Classification Result of Our Baseline

In this study, 96 REM data were taken from all three classes, a total of 288 REM period data were used. These data were cut into segments of 1s, 5s, 10s, and 20s in length. For comparison, expected results (chance level) are given for classification based on random voting [34].

The performance of classification based on kNN, SVM(RBF), LightGBM, GBDT, and RF. This is shown in Table 4, and the parameter of classifiers in Table 5. From a time period perspective, the 1s data segment has the lowest accuracy rate, and the 20s data segment has the highest accuracy rate. For example, the binary accuracy of CSP feature in LightGBM classifiers of 1s, 5s, 10s and 20s are 0.7150, 0.7926, 0.8435 and 0.8738 respectively. The same pattern is also seen in the other classification processes, which may be because some subjects received noise interference of 1s to 2s per minute during EEG acquisition process. From the perspective of features, CSP has higher accuracy than DE and PSD in binary classification and slightly less accuracy in multi-classification. For instance, the binary LightGBM accuracy of different features of 10s data are 0.8230, 0.8216 and 0.8435 for PSD, DE and CSP data respectively, while the accuracy of multi classification are 0.7568, 0.7587 and 0.7440 respectively. This indicates that the spatial-frequency domain (CSP) features have better classification accuracy than the frequency domain features (PSD, DE) for negative and positive classification, and similar findings were obtained in the study by Koelstra et al.[35].

**Table 4.** Classification accuracy based on 5 classifiers in different window sizes of PSD, DE, CSP and chance level

| Feature | Classifier | 1s |  | 5s |  | 10s |  | 20s |  |
| --- | --- | --- | --- | --- | --- | --- | --- | --- | --- |
|  |  | binary | multi | binary | multi | binary | multi | binary | multi |
| PSD | kNN | 0.5961 | 0.4597 | 0.7044 | 0.5909 | 0.7413 | 0.6298 | 0.7401 | 0.6502 |
|  | SVM | 0.6584 | 0.5196 | 0.7539 | 0.6512 | 0.7700 | 0.6979 | 0.7771 | 0.7160 |
|  | LightGBM | 0.6755 | 0.5357 | 0.7793 | 0.6828 | 0.8230 | 0.7568 | 0.8483 | 0.8149 |
|  | GBDT | 0.6803 | 0.5408 | 0.7790 | 0.6919 | 0.8238 | 0.7618 | 0.8502 | 0.8013 |
|  | RF | 0.6821 | 0.5385 | 0.7779 | 0.6761 | 0.8116 | 0.7415 | 0.8382 | 0.7857 |
| DE | kNN | 0.6530 | 0.5192 | 0.7402 | 0.6284 | 0.7638 | 0.6878 | 0.8220 | 0.7605 |
|  | SVM | 0.6823 | 0.5522 | 0.7754 | 0.6778 | 0.8063 | 0.7412 | 0.8428 | 0.8049 |
|  | LightGBM | 0.6995 | 0.5681 | 0.7859 | 0.6943 | 0.8216 | 0.7587 | 0.8553 | 0.8199 |
|  | GBDT | 0.7012 | 0.5707 | 0.7867 | 0.6935 | 0.8236 | 0.7594 | 0.8617 | 0.8179 |
|  | RF | 0.7030 | 0.5642 | 0.7755 | 0.6761 | 0.7957 | 0.7308 | 0.8280 | 0.7879 |
| CSP | kNN | 0.6866 | 0.5335 | 0.7635 | 0.6289 | 0.8149 | 0.6933 | 0.8572 | 0.7442 |
|  | SVM | 0.6835 | 0.5333 | 0.7628 | 0.6370 | 0.8220 | 0.7076 | 0.8636 | 0.7674 |
|  | LightGBM | 0.7150 | 0.5672 | 0.7926 | 0.6751 | 0.8435 | 0.7440 | 0.8738 | 0.7896 |
|  | GBDT | 0.7147 | 0.5694 | 0.7931 | 0.6805 | 0.8449 | 0.7430 | 0.8812 | 0.7967 |
|  | RF | 0.7177 | 0.5713 | 0.7954 | 0.6778 | 0.8419 | 0.7397 | 0.8757 | 0.7873 |
| Chance Level | kNN | 0.5034 | 0.3426 | 0.5069 | 0.3550 | 0.4870 | 0.3572 | 0.4953 | 0.3603 |
|  | SVM | 0.5006 | 0.3379 | 0.4931 | 0.3316 | 0.4991 | 0.3162 | 0.4768 | 0.3146 |
|  | LightGBM | 0.5051 | 0.3433 | 0.5046 | 0.3300 | 0.4861 | 0.3220 | 0.4861 | 0.3185 |
|  | GBDT | 0.5094 | 0.3426 | 0.5079 | 0.3271 | 0.4667 | 0.3325 | 0.4675 | 0.3198 |
|  | RF | 0.5100 | 0.3468 | 0.5051 | 0.3395 | 0.4889 | 0.3012 | 0.4675 | 0.3564 |

**Table 5.** Various parameters of classifiers

| Classifier | kNN | SVM | LightGBM | GBDT | RF |
| --- | --- | --- | --- | --- | --- |
| Parameters | n_neighbors=10 | C=8<br>gamma=0.25 | n_estimators=1000 | n_estimators=1000<br>max_depth= 8<br>max_features = 4 | n_estimators=1000 |

As the CSP features give the best classification results, the classification results of the CSP features based on random voting are given as a comparison. The mean of the accuracy of random voting for these classifiers of total segments, classifiers and features was 0.4946 for the binary classification and 0.3348 for the multi-classification, which is in line with the random voting predictions. This illustrates that the DEED dataset is effective in terms of classification performance and does not suffer from the influence of the classifiers. In summary, DEED dataset can be used for research on the classification of dream emotions.

## 7 Conclusion and Future Work

In this paper, we designed an experiment to collect dream EEG segments during the REM period and corresponding emotional state of the dream. The correlation between the subject’s subjective ratings of their emotions and the EEG was then analysed, showing a significant correlation in the alpha wave (p<0.01), which suggests a high correlation between alpha waves and dream emotions. Unsupervised clustering analysis was also performed on brain network patterns, and the clustering results for all five frequency bands showed significant patterns. We also conducted a hemispheric asymmetry analysis and found differences in the left and right hemisphere asymmetry index analysis between positive and negative emotions, with the right hemisphere playing a more important role in dream emotions. Finally, we evaluated the classification performance of DEED dataset and found that the multi-band CSP features had a higher binary classification accuracy and similar multi classification compared to DE and PSD, and accuracy rises with the segment length. This means the good classification performance of our dataset with reasonable preprocessing and feature extraction. The dataset provides samples of time series signals containing dream emotion information with non-stationary characters, obscure spatiotemporal features and time-series signals, suitable for algorithms that can be used for further AI research, such as bio-intelligence performance, shot less learning, generalisation and interference immunity. If there is a further understanding of the dataset relative to this, we have also provided Video list.xls and Emotion rating.xls. Combined with the raw data, this may also be illuminating in exploring the neural manifestations of biological characteristics of dreams, the interactions mechanisms of sleep emotion, and transitions between the conscious and unconscious.

It is undeniable that human brain function can be affected by the age, mental state and illness of the subject, and in this paper, only experimental data from healthy young adults were collected. Therefore, we should increase the sample size of subjects in subsequent experiments and analyse the dream emotions generated by different ages, occupations and environments in order to draw more profound conclusions.

